# NUCLEAR FACTOR Y, subunit A (NF-YA) proteins positively regulate flowering and act through *FLOWERING LOCUS T*

**DOI:** 10.1101/066399

**Authors:** Chamindika L. Siriwardana, Nerina Gnesutta, Roderick W. Kumimoto, Daniel S. Jones, Zachary A. Myers, Roberto Mantovani, Ben F Holt

**Affiliations:** Department of Microbiology and Plant Biology, University of Oklahoma, Norman OK, USA; Departimento di BioScienze, 6, Milan, Italy; Department of Plant Biology, University of California, Davis CA, USA

## Abstract

Photoperiod dependent flowering is one of several mechanisms used by plants to initiate the developmental transition from vegetative growth to reproductive growth. The NUCLEAR FACTOR Y (NF-Y) transcription factors are heterotrimeric complexes composed of NF-YA and histone-fold domain (HFD) containing NF-YB/NF-YC, that initiate photoperiod-dependent flowering by cooperatively interacting with CONSTANS (CO) to drive the expression of *FLOWERING LOCUS T (FT)*. This involves NF-Y and CO binding at distal *CCAAT* and proximal “CORE” elements, respectively, in the *FT* promoter. While this is well established for the HFD subunits, there remains some question over the potential role of NF-YA as either positive or negative regulators of this process. Here we provide strong support, in the form of genetic and biochemical analyses, that NF-YA, in complex with NF-YB/NF-YC proteins, can directly bind the distal *CCAAT* box in the *FT* promoter and are positive regulators of flowering in an *FT*-dependent manner.

**Author Summary:** For plants to have reproductive success, they must time their flowering with the most beneficial biotic and abiotic environmental conditions - after all, reproductive success would likely be low if flowers developed when pollinators were not present or freezing temperatures were on the horizon. Proper timing mechanisms for flowering vary significantly between different species, but can be connected to a variety of environmental cues, including water availability, temperature, and day length. Numerous labs have studied the molecular aspects of these timing mechanisms and discovered that many of these pathways converge on the gene *FLOWERING LOCUS T (FT)*. This means that understanding precisely how this gene is regulated can teach us a lot about many plant species in both natural and agricultural settings. In the current study, we focus on day length as an essential cue for flowering in the plant species *Arabidopsis thaliana*. We further unravel the complexity of *FT* regulation by clarifying the roles of *NUCLEAR FACTOR Y* genes in day length perception.

## Introduction

Plants undergo numerous developmental phase changes that are both species specific and intimately linked to the environments in which they evolved. One of the most important phase changes - as evidenced by the numerous pathways controlling the process – is the transition from vegetative to reproductive growth (recently reviewed in (1)). For many plant species, a potent trigger of the transition to reproductive growth is photoperiod-dependent flowering. Photoperiod-dependent species use the relative length of day and night to either activate or repress flowering such that it is timed with the appropriate environmental conditions to maximize reproductive success.

The model plant *Arabidopsis thaliana* (Arabidopsis) is a so-called long day plant; that is, it flowers rapidly when days are longer than ~12 hrs (2–5). Central to measuring photoperiod is the circadian regulation of *CONSTANS* (*CO*) transcription and the light-mediated regulation of CO protein accumulation (6). CO protein is stabilized in long days and is able to bind and transcriptionally activate *FLOWERING LOCUS T* (*FT*) (7, 8). FT protein is the principal mobile hormone - or “florigen” - that travels from leaves, where the photoperiod signal is perceived, to the shoot apex, where the floral transition occurs (9–12). In the shoot apex FT activates its downstream targets, which includes *SUPPRESSOR OF CONSTANS 1* (*SOC1*) and *APETALA 1* (*AP1*). Members of the heterotrimeric NUCLEAR FACTOR-Y (NF-Y) transcription factor family are required for activation of the *FT* promoter, thus initiating the downstream events leading to the floral transition (13–18).

NF-Y transcription factors are composed of three independent protein families, NF-YA, NF-YB, and NF-YC. To activate target genes, NF-YB and NF-YC dimerize in the cytoplasm and move to the nucleus where the heterodimer interacts with NF-YA to create the DNA-binding, heterotrimeric NF-Y transcription factor (21–24). NF-Y binding is widely regarded as sequence specific to the evolutionarily conserved *CCAAT* motifs, with some modified sites having been reported (15, 25, 26). All direct contacts with the pentanucleotide are made by NF-YA, while the NF-YB/NF-YC dimer primarily makes non-sequence specific contacts in adjacent regions, stabilizing the complex (27). NF-Y subunits have undergone an extensive expansion in plants (19, 20). For example, Arabidopsis has ten members of each *NF-Y* gene family (20).

Several NF-YB and NF-YC subunits have been demonstrated to regulate photoperiod dependent flowering (13, 16–18, 28, 29). Briefly, *nf-yb2 nf-yb3* double and *nf-yc3 nf-yc4 nf-yc9* triple mutants flower very late under normally inductive photoperiods (17). In both cases, the single mutants have either no effect or comparatively mild effects on flowering time, indicating overlapping functions for these family members. NF-YB and NF-YC proteins can physically interact with CO and loss of function mutations lead to *FT* expression downregulation (13, 16–18, 28). Finally, genetic and biochemical data suggest that NF-Y complexes bind the *FT* promoter at a distal *CCAAT* box (−5.3kb from start codon), while CO binds several clustered proximal **CO** regulatory elements (CORE – approx. −200bp upstream from start). Chromatin loops may stabilize the interactions between these two distally separated, DNA-bound complexes (8, 14, 30, 31).

In light of the NF-Y HFD interactions with CO in photoperiod-dependent flowering, immediate questions are whether NF-YA proteins are regulators of photoperiod-dependent flowering and whether this is CO-dependent and exerted through regulation of *FT*. Related to NF-YA roles in flowering, initial reports demonstrated that they can negatively regulate flowering as overexpression of some *NF-YA* genes caused late flowering (18, 32). Because NF-YA and CO proteins share a region of sequence homology, one possibility is that they compete for occupancy on NF-YB/C dimers: in this scenario, NF-YA and CO might play opposing negative and positive roles, respectively. However, recent reports suggest a more complex scenario, given 1) Genetic evidence for the importance of the −5.3kb *FT CCAAT* box in flowering (14, 30, 31); 2) DNA bound mammalian NF-Y crystal structure showing that NF-YA makes the direct contacts with the *CCAAT* box and that CO shows differences in amino acids necessary for these contacts (14, 27, 33, 34); and 3) Evidence that CO directly binds CORE sites (8). In addition, Hou et al. (15) suggested that NF-YA2 was a positive regulator of flowering time, but, surprisingly, that this was mediated by interaction with a novel, non-CCAAT cis regulatory element called NF-YBE in the *SOC1* promoter, and not the binding and regulation of *FT* expression.

As reported for *co* mutants (35), multiple groups have demonstrated that *nf-yb* and *nf-yc* mutants also had strongly reduced *FT* expression and that these reductions were directly correlated with alterations in flowering time (16–18, 28, 36). Likewise, overexpression of *NF-YB* and *NF-YC* genes was associated with *FT* upregulation (16, 28, 37–39). Mutations in cis-regulatory elements bound by either CO or NF-Y complexes in the *FT* promoter (*CCAAT* and/or CORE, respectively) also reduced *FT* expression in a manner that was directly correlated with the severity of flowering delays (14, 16, 17, 30). Further, constitutive overexpression of *CO* drove increased *FT* expression and early flowering, but these phenotypes were strongly reduced or eliminated in *nf-yb* and *nf-yc* mutants or when the −5.3kb *CCAAT* site was eliminated (17, 30, 38). Finally, multiple labs have shown *in vivo* and *in vitro* binding of NF-Y and CO proteins to the *FT* promoter and mutations in the associated *CCAAT* and CORE cis-regulatory elements additively reduce *FT* expression and delay flowering (7, 8, 14, 30). Thus, it remains very well-supported that photoperiod-dependent flowering is mediated through direct regulation of *FT* by CO and NF-Y complexes.

Here we address the roles of NF-YA proteins in *FT* binding, expression regulation, and photoperiod-dependent flowering time. Using a combination of genetic and biochemical approaches, we show complete NF-Y complexes, including NF-YA, bound to the −5.3kb *FT CCAAT* box. We further demonstrate that NF-YA and NF-YB constructs that can drive early flowering do this activity in an FT-dependent manner. Because *SOC1* is downstream of *FT* (40), our data further indicate that *FT* is a key regulatory target of NF-Y/CO complexes in the photoperiod-dependent flowering pathway.

## Results

### *NF-YA* genes can be positive regulators of photoperiod dependent flowering

To identify NF-YAs involved in flowering, we first examined constitutive overexpression (35S promoter) in first generation (T1) transgenic plant lines for each of the 10 Arabidopsis *NF-YA* genes (lines described in (41). We observed that *p35S:NF-YA2* and *p35S:NF-YA6* expressing plants consistently flowered earlier than Col-0. Nevertheless, confident interpretations of these data were complicated by the pleiotropic, dwarf phenotypes in most overexpressing lines. In fact, lines that constitutively overexpressed *NF-YA6* were infertile and did not survive (as previously described, (41)). We were able to isolate and quantify stable, third generation transgenic *p35S:NF-YA2* lines and compare them to several other stable lines for constitutively expressed *NF-YA* genes (Fig 1A). Two independent *p35S:NF-YA2* lines flowered early (~10 leaves, compared to 13 for wild type Col-0 plants), while overexpression of other *NF-YA* genes either did not alter flowering or actually caused modestly later flowering. This is consistent with the original observations of Wenkel (18). We note that all of these plant lines showed similar dwarf phenotypes, suggesting that our flowering time observations were not directly correlated with this phenotype.

**Fig 1.**
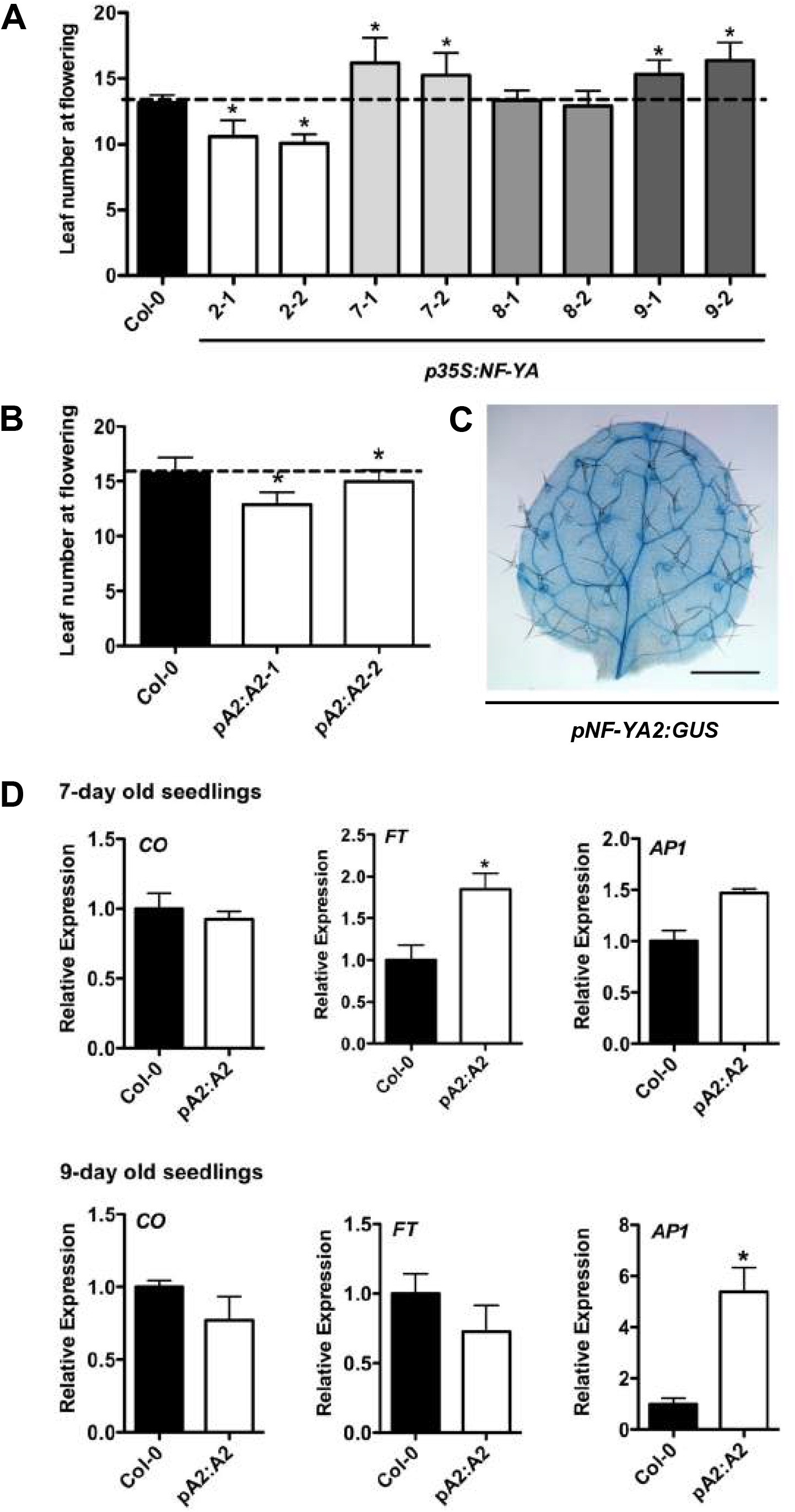
NF-YA2 is a positive regulator of photoperiod dependent flowering. A) Flowering time quantification of two independent *p35S:NF-YA2, p35S:NF-YA7, p35S:NF-YA8*, and *p35S:NF-YA9* plant lines. B) Flowering time quantification of two independent *pNF-YA2:NF-YA2* plant lines. C) The expression pattern of *pNF-YA2-GUS* in leaves of 10 day old plants. D) Expression of *CO, FT*, and *AP1*. Asterisks in 1A and 1B represent significant differences derived from one-way ANOVA (P < 0.05) followed by Dunnett’s multiple comparison post hoc tests against Col-0. Asterisks in 1D represent significant differences derived from Student’s T-tests (p<0.05).

To avoid the pleiotropic effects from ectopically overexpressing *NF-YA2*, we additionally generated stable, native promoter transgenic plant lines (*pA2:NF-YA2*). Presumably due to position effects, some of these lines expressed high levels of *NF-YA2* (~60 fold overexpressed) and were early flowering (Fig 1B, S1 Fig). Interestingly, these plants appeared phenotypically normal, suggesting that the dwarf phenotypes of p35S-driven lines is more related to ectopic expression than overexpression, *per se*. Note that our previous research on *NF-Y:GUS* expression patterns showed that both *NF-YA2* and *NF-YA6* had very strong vascular expression, consistent with the expected localization of floral promoting genes (Fig 1C and (17, 30, 31, 33, 42, 43).

As discussed above, previous reports suggest that CO, NF-YB and NF-YC regulate flowering primarily by controlling *FT* expression which, in turn, rapidly upregulates *AP1* (16, 17, 28, 30, 31, 35, 40, 44). We used the stable *pNF-YA2:NF-YA2-1* plant line to test if *NF-YA2* regulates the same set of genes. We used the time points of seven and nine days after germination because they correlate with the initiation of flowering signals in long day grown plants (42). *NF-YB* and *NF-YC* do not affect the expression of *CO* (16, 17, 28); likewise, *CO* was not misregulated in the *NF-YA2* overexpressor (Fig 1C). However, the expression of *FT* was upregulated in seven day old *pNF-YA2:NF-YA2* plants, which was followed by significant *AP1* upregulation by day nine. These results suggest that *NF-YA2*, like its *NF-YB* and *NF-YC* counterparts, regulates flowering by controlling *FT* expression.

### The NF-YB2^E65R^ mutation prevents NF-YA subunits from entering into NF-complexes

Because of the apparent difficulties in working directly with NF-YAs, the likely overlapping functionality between family members in flowering (e.g., Hou reports that *nf-ya2* mutants have no flowering delay, (15)), and lethality (45, 46), we decided to indirectly manipulate NF-YA function by altering its ability to interact with the HFD dimer. In mammals, the NF-YB^E92R^ mutant protein specifically loses interaction with NF-YA, but not NF-YC (22). Crystal structure analysis of the NF-Y complex demonstrated that this glutamic acid makes multiple contacts with NF-YA Arg249 and Arg253 (27). Alignments between human and Arabidopsis NF-YB proteins show that this glutamic acid (E65 in Arabidopsis NF-YB2) is completely conserved (Fig 2A) and examination of other published alignments also confirm this conservation in the monocot lineage (33, 47–49). Thus, we reasoned that NF-YB2^E65R^ mutations would eliminate the ability of NF-YA to enter floral promoting NF-Y complexes and allow us to further test the hypothesis that NF-YA proteins are positive regulators of photoperiod-dependent flowering.

**Fig 2.**
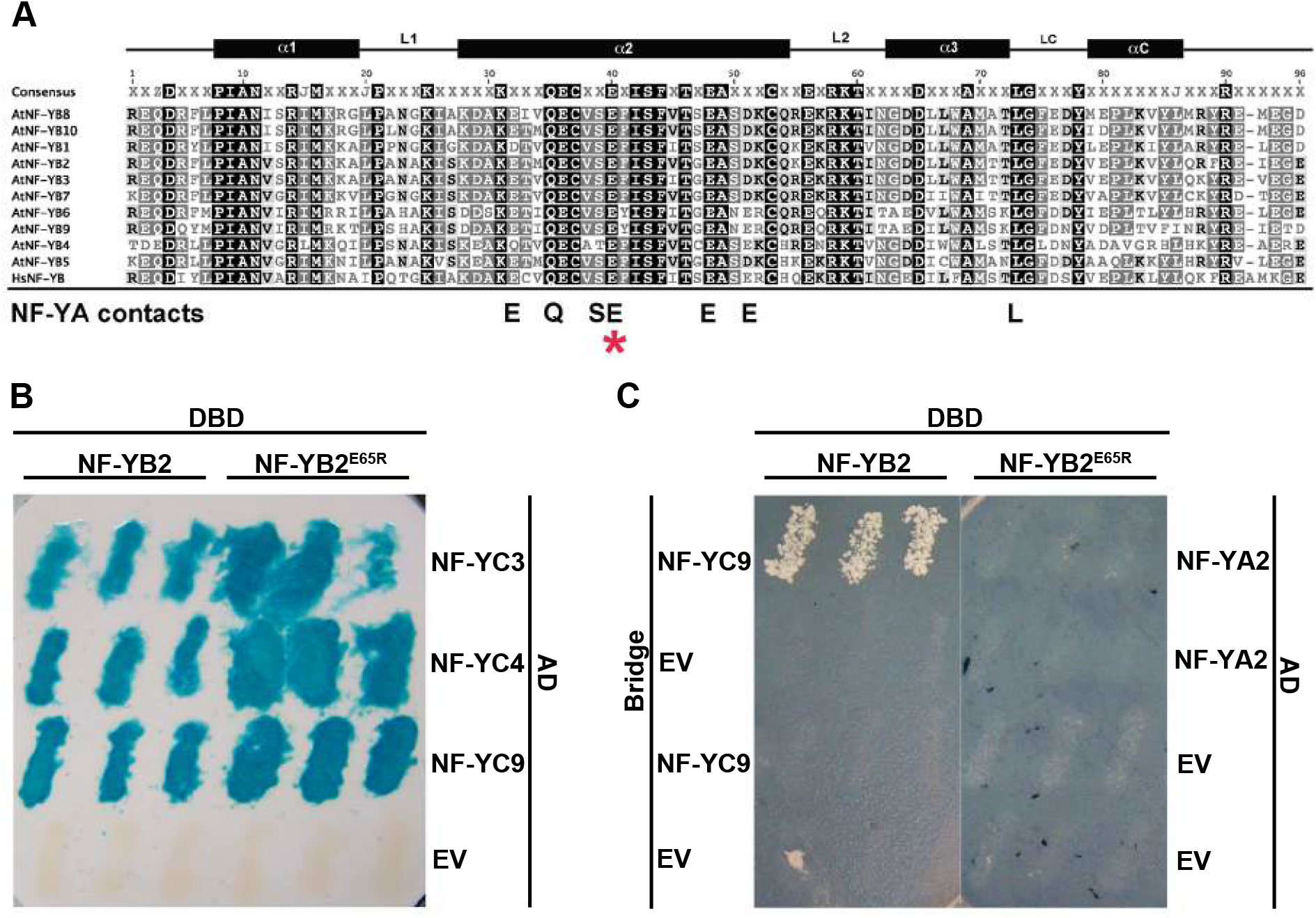
NF-YB2^E65R^ loses interaction with NF-YA subunits. A) Alignment of the core domain of human and Arabidopsis NF-YB subunits. * marks the position of the conserved glutamic acid required for interaction with NF-YA in humans (27). B) NF-YB2 and NF-YB2^E65R^ interact with NF-YC3, NF-YC4, and NF-YC9 in Y2H assays. C) NF-YB2, but not NF-YB2^E65R^, interacts with NF-YA2 when NF-YC9 is expressed using a bridge vector in yeast three-hybrid assays.

We first used yeast two hybrid assays to test if NF-YB2^E65R^ could interact with NF-YC3, NF-YC4, and NF-YC9: indeed, we found that both NF-YB2 and NF-YB2^E65R^ were able to physically interact with the NF-YCs (Fig 2B). Since NF-YA trimerizes with HFD dimers and not individually with NF-YB or NF-YC (50), we used yeast three hybrid assays to test the ability of NF-YA2 to enter into a complex with NF-YB2^E65R^ and NF-YC9 (Fig 2C). As predicted, NF-YA2/NF-YB2/NF-YC9 complexes formed, but the NF-YB2^E65R^ variant prevented formation of the trimeric NF-Y complex. Thus, the NF-YB2^E65R^ provides a powerful genetic tool to test the requirement for NF-YA in photoperiod-dependent flowering.

### The NF-YB2^E65R^ mutation prevents rescue of a late flowering *nf-yb2 nf-yb3* mutant

We predicted that *p35S:NF-YB2^E65R^* would be unable to drive early flowering in wild type Col-0 or rescue the *nf-yb2 nf-yb3* late flowering phenotype. We tested this by overexpressing both *p35S:NF-YB2* and *p35S:NF-YB2^E65R^* in each background. As previously described, here and throughout this study, we examined T1 plants as it gave a better representation of the response by eliminating bias associated with the selection of individual transgenes. For each transgene we examined 15–20 individual plants and for selected experiments we generated two independent T3 transgenic lines for further testing (14). We found that *p35S:NF-YB2* showed a trend towards earlier flowering in Col-0, but only caused significantly earlier flowering in a subset of independent experiments (Fig 3A, non-significant example shown). However, *p35S:NF-YB2 nf-yb2 nf-yb3* plants flowered ~20 leaves earlier than the parental mutant (Fig 3B–D). With the *p35S:NF-YB2^E65R^* version, Col-0 actually flowered significantly later than normal (indicating dominant interference with the endogenous complexes) and there was no rescue of the *nf-yb2 nf-yb3* late flowering phenotype (Fig 3A–D).

**Fig 3.**
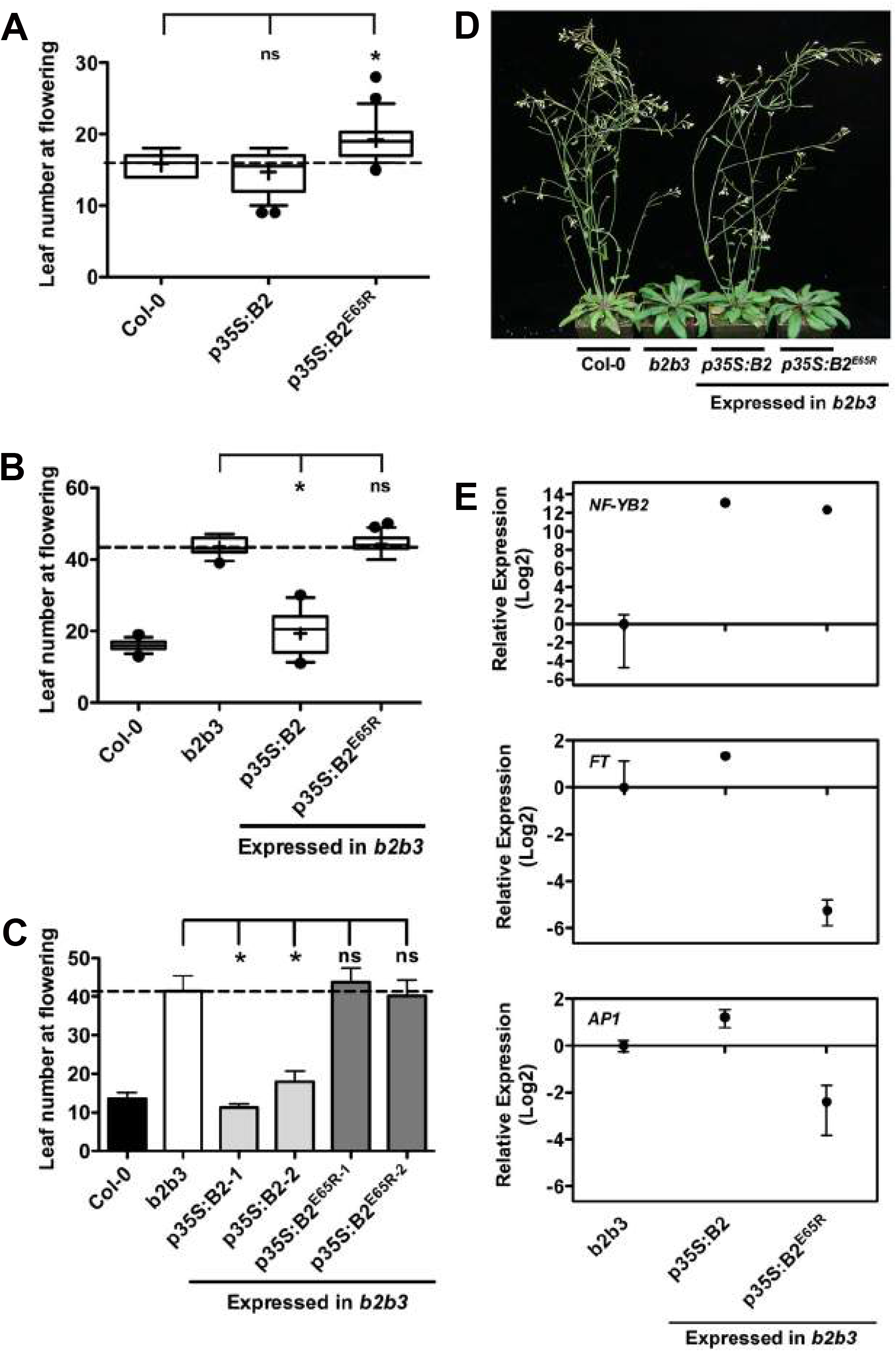
*p35S:NF-YB2^E65R^* cannot rescue the *nf-yb2 nf-yb3* late flowering phenotype. A) Flowering time quantification of T1 *p35S:NF-YB2* and *p35S:NF-YB2^E65R^* plants in the Col-0 background. B) Flowering time quantification of T1 *p35S:NF-YB2* and *p35S:NF-YB2^E65R^* plants in the *nf-yb2 nf-yb3* background. C) Flowering time quantification of stable T3 *p35S:NF-YB2* and *p35S:NF-YB2^E65R^* plants in the *nf-yb2 nf-yb3* background. D) Representative plants of *p35S:NF-YB2* and *p35S:NF-YB2^E65R^* in the *nf-yb2 nf-yb3* background. E) Expression of *NF-YB2, FT* and *AP1* in the *nf-yb2 nf-yb3* background. Asterisks in 3A, 3B and 3C represent significant differences derived from one-way ANOVA (P<0.05) followed by Dunnett’s multiple comparison post hoc tests against *nf-yb2 nf-yb3*.

To confirm that NF-YB^E65R^ was localizing properly, we compared plants expressing *NF-YB2-YFP* and *NF-YB2^E65R^-YFP* and found that both had identical nuclear localization patterns (S2A Fig). Additionally, we measured NF-YB protein accumulation in late flowering *p35S:NF-YB2^E65R^* T1 plants (all >31 leaves at flowering) compared to a well-characterized, stable, early flowering *p35S:NF-YB2* line (all proteins were translationally fused to the HA epitope). The *p35S:NF-YB2^E65R^* T1 lines showed the expected variation in NF-YB protein accumulation; note that even lines that strongly accumulated NF-YB2^E65R^ could not rescue late flowering (S2B Fig; e.g., compare protein accumulation in *p35S:NF-YB2^E65R^* lines 6, 10, 11, and 12 to the stable p35S:NF-YB2 line). Stable, single insertion T3 lines showed the same pattern of late flowering regardless of high NF-YB2^E65R^ accumulation (Fig 3C and S2C Fig). Finally, we compared stable *p35S:NF-YB2 nf-yb2 nf-yb3* and *p35S:NF-YB2^E65R^ nf-yb2 nf-yb3* for expression of *NF-YB2, FT*, and *AP1* (Fig 3E). Although both lines had very high, ~equivalent *NF-YB2* expression, *p35S:NF-YB2* resulted in increased *FT* and *AP1* expression while *p35S:NF-YB2^E65R^* significantly suppressed both. Collectively, we take these data as strongly suggestive data that NF-YA participation in trimer formation is important for the promotion of flowering.

### NF-YA2 and NF-YA6 heterotrimerize with NF-YB2 and NF-YC3 *in vitro* to bind the −5.3kb *CCAAT* box

We previously showed that NF-YB2 and NF-YC3, together with mouse NF-YA, are able to bind a 31bp, CCAAT-containing oligonucleotide from the *FT* −5.3kb site (14). At that time we were unsure of the likely Arabidopsis NF-YA(s) involved in flowering: with the data presented here and a recent publication (15) showing that NF-YA2 and NF-YA6 can act as positive regulators of flowering, we used EMSA to test if NF-YA2 and NF-YA6 are able to bind a probe encompassing the −5.3kb *CCAAT* box on *FT*. In the presence of NF-YB2/NF-YC3 dimers, NF-YA2 and NF-YA6 bound the *CCAAT* probe in a concentration-specific manner (Fig 4). However, neither NF-YA2 nor NF-YA6 could individually bind the *CCAAT* probe. Further, CO did not bind the *CCAAT* probe, individually or in the presence of the NF-YB2/NF-YC3 dimer. We additionally tested equivalent concentrations of NF-YA2 with the NF-YB2^E65R^/NF-YC3 and found that this combination completely lost the ability to bind the *CCAAT* probe. Collectively, this data shows that plant NF-Y complexes interact and bind the *FT* −5.3kb *CCAAT* box in a manner that is similar, if not identical, to the mammalian counterparts.

**Fig 4.**
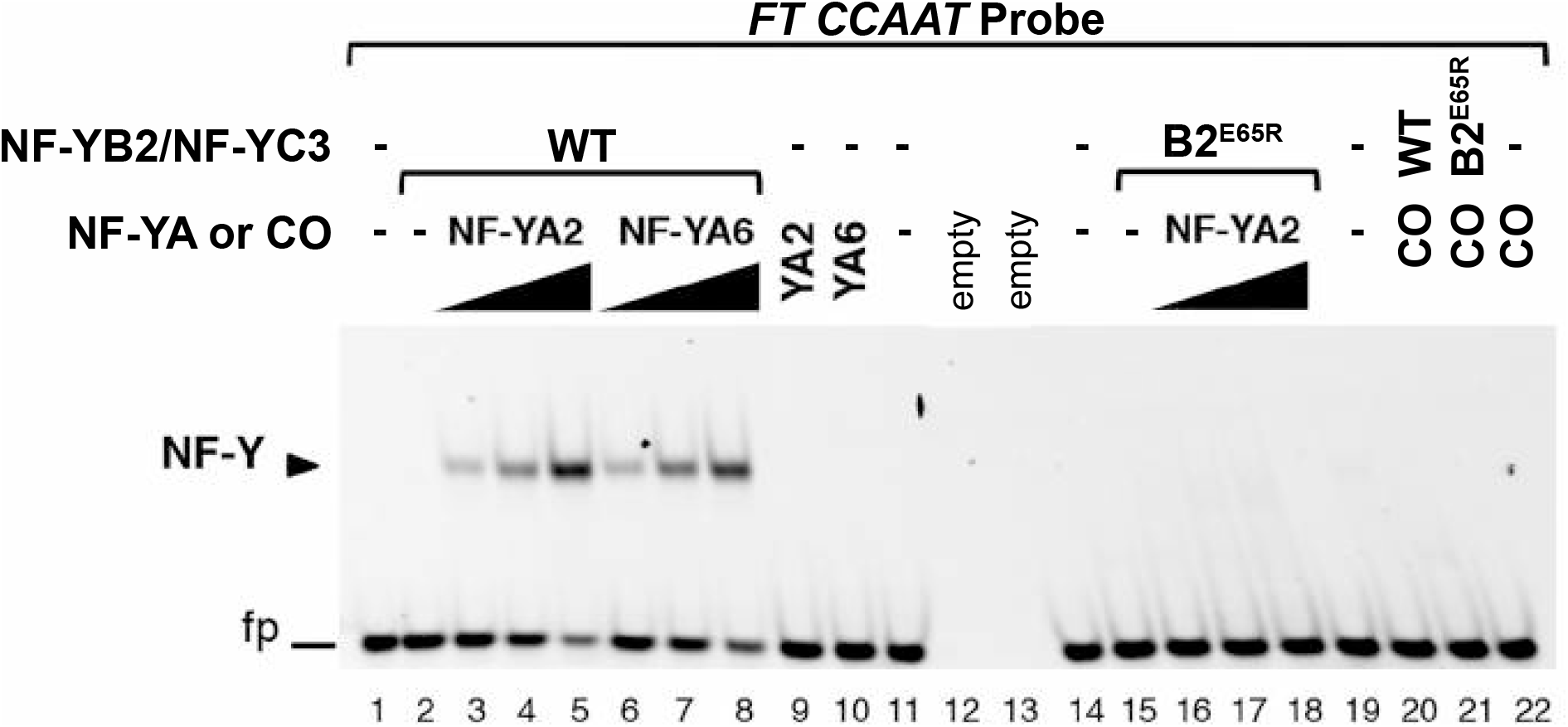
NF-YA2 and NF-YA6 bind the *FT* −5.3kb *CCAAT* box as a trimer with NF-YB2 and NF-YC3. NF-Y trimerization and *FT CCAAT* binding was assessed by EMSA analysis. An *FT CCAAT* probe was incubated with wild type (WT, lanes 2–8; 20) or E65R mutant (B2^E65R^, lanes 15–18; 21) NF-YB2/NF-YC3 dimers (60 nM) in the presence of NF-YA2 (lanes 3–5; 16–18), or NF-YA6 (lanes 6–8) at increasing molar ratios (3, 4.5 or 6 fold), or CO (lanes 20, 21; 6 fold molar ratio). As controls, NF-YA2 (lane 9), NF-YA6 (lane 10), or CO (lane 22) were incubated alone with the probe, at the highest concentration of the dose curve (360 nM), in the absence of NF-YB2/NF-YC3. Lanes 1, 11, 14, 19: probe alone, without protein additions; lanes 12, 13: empty lanes. The NF-Y/DNA complex is indicated by a labelled arrowhead. fp: free probe.

### *p35S:NF-YB2^E65R^* fused to a strong activation domain is not able to induce flowering in a CONSTANS-deficient mutant

A potential criticism of using NF-YB^E65R^ as a tool to demonstrate an NF-YA requirement in flowering is that we do not know how it might affect interactions with other components involved in photoperiod-dependent flowering. In particular, we do not know if it might impact CO recruitment or binding to its CORE site. However, when a strong transcriptional activation domain (called EDLL) was fused to NF-YB2, it was able to drive early flowering in a *co-9* loss of function mutant (38). Therefore, if NF-YA interactions are relevant in flowering, we expect that an NF-YB2^E65R^-EDLL would not be able to drive early flowering or rescue a *co* mutant.

We first overexpressed (35S) *NF-YB2-EDLL* in Col-0 and extended the findings to the *co-2* mutant in the Ler ecotype (Fig 5A–B): while *NF-YB2* alone did not drive early flowering, *NF-YB2-EDLL* expressing plants were consistently earlier, thus confirming previous data (38). However, in each case, *NF-YB2^E65R^-EDLL* either caused later flowering (presumably the dominant negative effect again, Fig 3A) or had no effect. We then used the *nf-yb2 nf-yb3* background where *NF-YB2* plants flowered at a mean of ~21 leaves and *NF-YB2-EDLL* flowered at ~12 leaves (Fig 5C); *NF-YB2^E65R^-EDLL* was once again unable to alter flowering time. Short day grown plants, which mimic a *co* mutant because CO is unable to accumulate (2), told the same story - *NF-YB2-EDLL*, but not *NF-YB2^E65R^-EDLL*, caused earlier flowering (Fig 5D). Finally, we repeated the entire transgenic panel in the loss of function *ft-10* mutant (Fig 5E). Importantly, all constructs, including *NF-YB2-EDLL*, failed to cause significantly earlier flowering in the *ft-10* genetic background. Collectively, this data adds additional evidence for NF-YA as a positive, FT-dependent regulator of photoperiod-dependent flowering.

**Fig 5.**
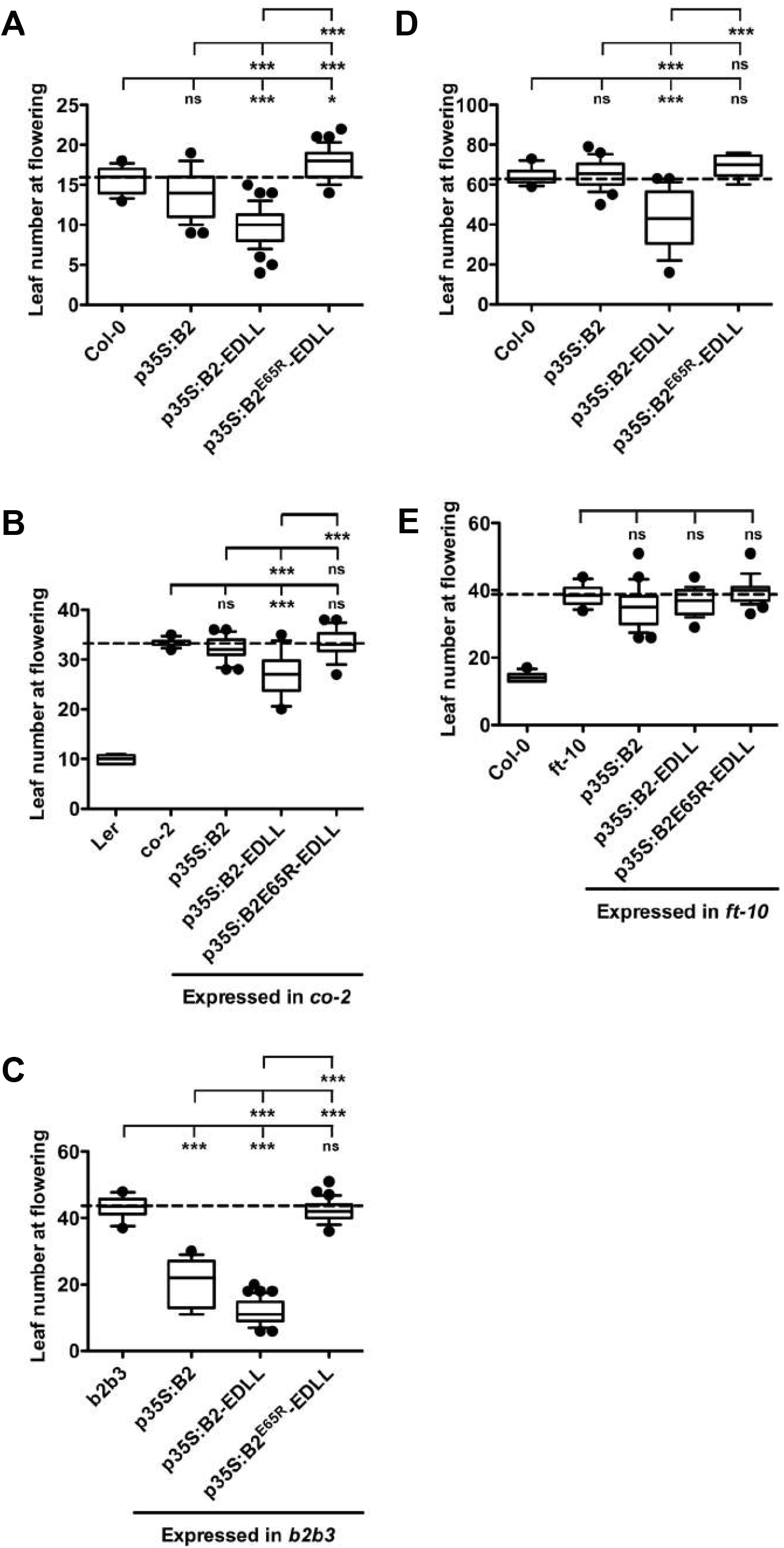
*NF-YB2-EDLL*, but not *NF-YB2^E65R^-EDLL*, rescues late flowering in an *FT*-dependent manner. T1 flowering time quantification of *p35S:NF-YB2, p35S:NF-YB2-EDLL*, and *p35S:NF-YB2^E65R^-EDLL* in A) Col-0 B) *co-2* C) *b2b3* D) short days E) *ft-10*. Asterisks represent significant differences derived from one-way ANOVA (P<0.05) followed by Bonferroni’s multiple comparison tests.

### *NF-YA2-EDLL* induces flowering in a CONSTANS-deficient mutant

We hypothesized that if NF-YA2 is able to interact with NF-YB/NF-YC dimers on the *FT* promoter, attaching the *EDLL* domain to the *pNF-YA2:NF-YA2* construct would also induce flowering in *co* mutants. If true, this would significantly extend the NF-YB2^E65R^ and EMSA results above, ameliorating possible concerns about relying on NF-YB2^E65R^ as a proxy measure of NF-YA function. Again, we first tested flowering responses in the Col-0 background. Both *pNF-YA2:NF-YA2* and *pNF-YA2:NF-YA2-EDLL* drove earlier flowering (Fig 6A). In the *co-2* background, *pNF-YA2:NF-YA2-EDLL* induced much earlier flowering (~20 leaves earlier than *co-2*), whereas the control *pNF-YA2:NF-YA2* did not (Fig 6B). As with *NF-YB2-EDLL* (Fig 5E), *NF-YA2-EDLL* was completely unable to induce flowering in the *ft-10* background (Fig 6C), indicating once again an *FT*-dependent, positive role for NF-YA proteins in flowering.

**Fig 6.**
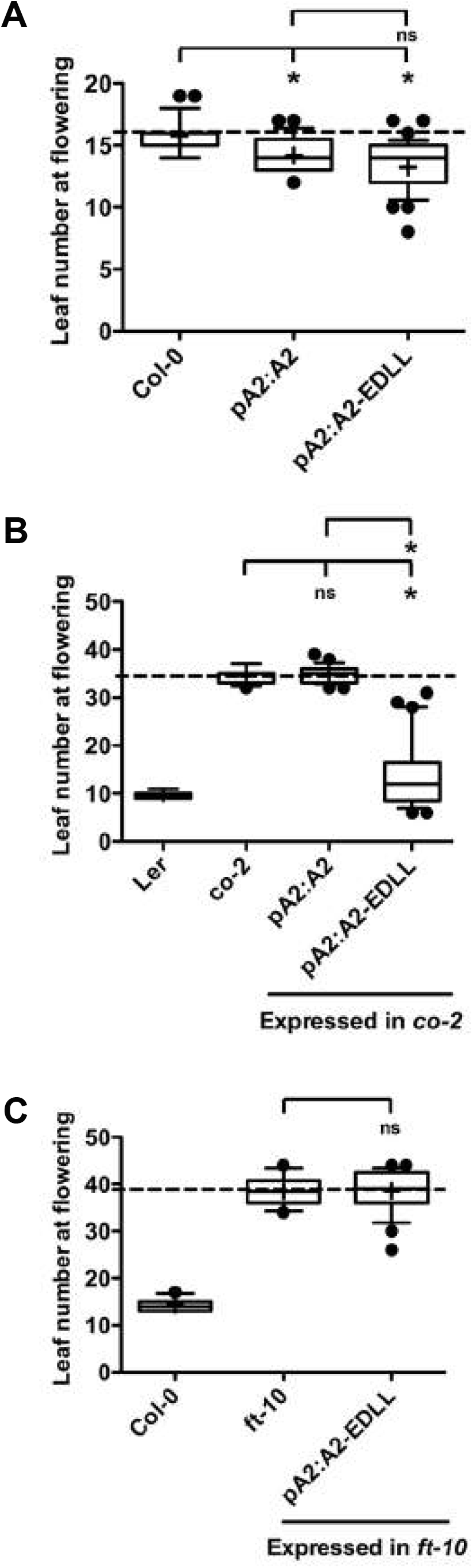
*pNF-YA2:NF-YA2-EDLL* can induce flowering in the absence of CO. Flowering time in A) Col-0, B) *co-2*, and C) *ft-10*. Asterisks represent significant differences derived from one-way ANOVA (P < 0.05) followed by Bonferroni’s multiple comparison tests.

## Discussion

Our initial understanding of NF-Y roles in flowering was primarily driven by evidence of physical interactions between individual NF-Y subunits and CO, as well as *in planta* overexpression analyses (13, 18). Thereafter, loss of function mutations in HFD subunits identified specific *NF-YB* and *NF-YC* genes involved in flowering (16, 28, 51). Demonstrating roles for *NF-YAs* has proven more difficult, since they appear to have redundant functions, and overexpressing them leads to substantially deleterious pleiotropic effects (41, 52, 53). Here we have attempted to work around these difficulties with a variety of biochemical and genetic approaches. We provide a compelling body of evidence that NF-YA2 and NF-YA6, and perhaps other NF-YAs, can activate *FT* expression, and are FT-dependent, positive regulators of flowering.

Previously, NF-YAs were believed to act as negative regulators of flowering, because overexpression of two *NF-YA* genes, *NF-YA1* and *NF-YA4*, led to later flowering (18). We noticed the same response with *NF-YA7* and *NF-YA9* overexpressors. In another study by Leyva-Gonzalez (52), this was also the outcome of generalized overexpression of *NF-YAs*. A recent publication showed that NF-YA2 represses stress-mediated flowering responses (32). Further *miR169* was shown to target and degrade *NF-YA2* transcripts, which led to an induction of flowering through the downregulation of *FLC* and resulting upregulation of *FT*. However, there were a few question areas that were not clearly addressed. Loss-of-function mutants of *FLC* do not have an effect on flowering in Col-0 plants under LD conditions (54), and how the down regulation of *FLC* led to the flowering phenotypes under these conditions is not clear. Nevertheless, our observation of early flowering in *NF-YA2* overexpression lines is consistent with those recently reported (15). The central role of *FT* in the regulation of flowering has been established, and the recent report that NF-Ys regulate photoperiod-dependent flowering *via SOC1*, instead of *FT* (15), seems at odds with existing evidence, as well as experiments presented here. Elegant genetic experiments previously demonstrated that *SOC1* activation is downstream of *FT* in a linear pathway (40). Therefore, if NF-Ys are directly binding and activating *SOC1* to activate photoperiod-dependent flowering, *FT* loss of function alleles (such as *ft-10* used here) should not impair this function. However, we find that *p35S:NF-YA-EDLL* and *p35S:NF-YB-EDLL* cannot drive early flowering in the absence of *FT*, strongly suggesting that *SOC1* is not their only target in photoperiod-dependent flowering. We do not rule out the possibility that the NF-Y are also involved in the direct regulation of *SOC1;* however, regulation of *SOC1* alone cannot explain the flowering phenotypes discussed here.

Regulation of the *FT* promoter is influenced by a plethora of pathways and numerous cis-regulatory elements continue to emerge (14, 30, 31, 55). One of these is the −5.3kb *CCAAT* enhancer site, where both deletions and mutations significantly delay flowering time (14, 30, 56). We provide here formal *in vitro* evidence that complexes formed by NF-YA2 and NF-YA6, associated with NF-YB2/NF-YC3, robustly and specifically bind to this site. Interestingly, the phenotype of the −5.3kb *CCAAT* mutant was not as strong as those from *nf-y* HFD loss of function alleles (14, 16), implying that there must be additional *CCAAT* sites bound by the NF-Y trimer in the *FT* promoter or that NF-Y subunits also regulate non-CCAAT sites. Another set of important sites responsible for CO activation, CORE, are in the proximal promoter (8, 30). Indeed, the near complete loss of photoperiod-dependent flowering responses in *nf-yb2 nf-yb3* and *co* mutants strongly argues that NF-Y complexes and CO must be necessary for function at both cis-regulatory regions. In keeping with this, we recently showed that NF-Y, bound to the −5.3kb *CCAAT*, and CO, bound to CORE sites, physically interact via a chromatin loop. Further, simultaneous mutations in the −5.3kb *CCAAT*, CORE1 and CORE2 sites in the *FT* promoter nearly eliminated rescue of an *ft-10* mutant (14). The importance of the −5.3 kb *CCAAT* element implies a role of the sequence-specific subunit NF-YA; however, the interactions of the HFD subunits with CO, and the resulting enhancer-promoter connections through CORE, made the direct demonstration of NF-YA function in *FT* expression and flowering all the more important.

NF-YB^E65R^ overexpressors were not able to rescue the late flowering phenotype of the *nf-yb2 nf-yb3* mutant. We formally excluded that this was due to expression levels and we could also exclude that the mutant folded incorrectly for two reasons: 1) Recombinant production in *E. coli* recovered wt and E65R as soluble proteins when co-expressed with NF-YC3, and indeed both were easily purified, and 2) The mutant had a dominant negative effect on flowering time when overexpressed in Col-0 plants. A similar conclusion on the dominant negative nature of the glutamic acid mutation was made for rat NF-YB (CBF-A) *in vitro* (22), but this is the first demonstration that it could also act *in vivo*. The likeliest explanation for the dominant negative behavior of NF-YB2^E65R^ is related to formation of HFD heterodimers impaired in trimer formation, and hence normal NF-Y function – i.e., it is possible that they subtract functional NF-YCs, which would otherwise enter the normal trimerization/CCAAT-binding processes. Obviously, we cannot formally rule out the possibility that the NF-YB2^E65R^ mutant lost interaction with proteins other than NF-YA and that this resulted in the lack of rescue of late flowering.

To rule out the possibility that the NF-YB2 flowering phenotypes were possibly due to loss of interaction with CO, we used the EDLL transactivation domain. CO was previously demonstrated to provide an activation domain for the NF-Y complex and NF-YB2 was able to drive flowering in the absence of CO when fused to the EDLL activation domain (38). However, in the current study, *p35S:NF-YB2^E65R^-EDLL* was not able to induce flowering indicating that while CO provides an activation domain for the NF-Y complex, the HFD dimer is non-functional in the absence of NF-YA. Our EMSA data further connects an NF-YA requirement to the capacity to bind at *CCAAT* elements. Finally, the flowering phenotypes for *pNF-YA2:NF-YA2-EDLL* were essentially the same as *p35S:NF-YB2-EDLL*. Both constructs were able to induce flowering in *co* mutants, were not able to induce flowering in *ft-10* mutants, and drove earlier flowering in Col-0. Collectively, these data strongly suggest that NF-YA2 is required for photoperiod-dependent flowering, acts directly on the *FT* promoter, and is FT-dependent.

## Methods

### Multiple sequence alignments

Protein sequences were obtained from TAIR (http://www.arabidopsis.org (57) or National Center for Biotechnology Information (http://www.ncbi.nlm.nih.gov/) and manipulated in TextWrangler (http://www.barebones.com) Multiple sequence alignments were made using ClustalX (58) and shaded within Geneious (http://www.geneious.com/).

### Generation of overexpression constructs

The *p35S:NF-YB2* and the ten *p35S:NF-YA* constructs were previously described (41, 49), as was the *35S* promoter (59). *NF-YB2^E65R^* was amplified from cDNA using mutagenic PCR. *pNF-YA2:NF-YA2* was amplified using genomic DNA with the promoter region starting approximately 1 KB upstream of the start codon. The proof reading enzyme Pfu Ultra II (cat#600670; Agilent Technologies) was used for PCR reactions and the resulting fragments were ligated into GATEWAY^™^ entry vector pENTR/D-TOPO (cat#45-0218; Invitrogen). The EDLL domain (38) was amplified from cDNA and contained *Acs1* sites, which were used to clone the EDLL domain into the pENTR/D-TOPO backbone of *NF-YB2* and *NF-YB2^E65R^* entry clones. All entry clones generated were sequenced and other than the point mutation were identical to sequences at TAIR (http://www.arabidopsis.org (57). Entry clones were sub-cloned into the following destination vectors using the GATEWAY™ LR Clonease II reaction kit (cat#56485; Invitrogen): *NF-YB2^E65R^* into pEarlyGate101 (60); *NF-YB2, NF-YB2-EDLL* and *NF-YB2^E65R^-EDLL* into pK7FWG2 (61) *pNF-YA2:NF-YA2* and *pNF-YA2:NF-YA2-EDLL* into pEarlyGate301 (60) S1 Table lists primer sequences used for cloning and mutagenesis.

### Plant transformation, cultivation and flowering time experiments

*Arabidopsis thaliana* ecotype Columbia (Col-0) was the wild type for all experiments. *nf-yb2 nf-yb3, ft-10 and co-2* (40, 49, 62) were previously described. Plants were transformed using Agrobacterium mediated floral dipping (63). Plants were cultivated in a custom-built walk-in chamber under standard long day conditions (16h light/8h dark) using plant growth conditions previously described (41). Leaf number at flowering was measured as the total number of rosette and cauline leaves on the primary axis at flowering.

### Protein expression and purification

The cDNAs encoding for NF-YA2 (aa 134-207) and NF-YA6 (aa 170-237) were obtained by gene synthesis (Eurofins Genomics) and cloned into pnEA/tH (64) by restriction ligation with NdeI and BamHI to obtain C-terminal 6His-tag fusions. The CCT domain of CONSTANS (aa 290-352), with the addition of a 5’ ATG, was cloned into pnEA/tH via PCR amplification followed by restriction ligation with XhoI and MunI to obtain C-terminal 6His-tag fusions. Clones were verified by sequence analysis. *NF-YB2* mutant cDNA, encoding for aa 24-116 with residue E65 mutated to R *(NF-YB2^E65R^)* was obtained by gene synthesis and subcloned in pET15b to obtain N-terminal 6His-tag fusion. 6His-NF-YB2 or 6His-NF-YB2^E65R^/NF-YC3 soluble HFD dimers were produced by co-expression in *E. coli* BL21(DE3) and purified by ion metal affinity chromatography (IMAC) as described in (65). NF-YA2-6His, NF-YA6-6His or CO-6His were expressed in BL21(DE3) by IPTG induction (0.4mM IPTG for 4h at 25C) and purified by IMAC (HisSelect, SIGMA-Aldrich) in buffer A (10mM Tris pH 8.0, 400mM NaCl, 2mM MgCl2, 5mM imidazole). Purified proteins were eluted in Buffer A containing 100mM imidazole, and dialysed against Buffer B (10mM Tris-Cl pH 8.0, 400mM NaCl, 2mM DTT, 10 % glycerol).

### Electrophoretic Mobility Shift Assays

EMSA analyses were performed essentially as previously described (14, 64, 65). Heterotrimer formation and CCAAT-box DNA-binding of wt or mutant NF-YB2/NF-YC3 dimers was assessed by addition of purified NF-YAs (or CO) using the Cy5-labeled *FT CCAAT* probe (14). DNA binding reactions (1μl) (20nm *FT CCAAT* probe, 12mM Tris-HCl pH 8.0, 50mM KCl, 62.5mM NaCl, 0.5mM EDTA, 5mM MgCl2, 2.5mM DTT, 0.2 mg/ml BSA, 5% glycerol, 6.25ng/μl poly dA-dT) were incubated with wt or mutant NF-YB2/NF-YC3 dimers (60nm), with or without NF-YA2 or-YA6 (or CO), as indicated in Figure 4. Proteins were pre-mixed in Buffer B containing 0.1 mg/ml BSA, then added to DNA binding mixes. After 30min incubation at 30C, binding reactions were loaded on 6% polyacrylamide gels and separated by electrophoresis in 0.25X TBE. Fluorescence gel images were obtained and analyzed with a Chemidoc^™^ MP system and ImageLab^TM^ software (Bio-Rad).

### Western Blot

Total protein was extracted by grinding in lysis buffer (20mM Tris, pH 8.0, 150mM NaCl, 1mM EDTA, pH 8.0, 1% Triton X-100, 1% SDS with fresh 5mM DTT, 10mM protease inhibitor). NF-YB2-YFP/HA and NF-YB2^E65R^-YFP/HA were detected using high affinity anti-HA primary antibody (cat#11 867 423 001; Roche) and goat anti-rat secondary antibody (cat#SC-2032; Santa Cruz Biotechnology). Horseradish peroxidase-based ECL plus reagent was used for visualization in a Bio-Rad ChemiDoc XRS imaging system. The membrane was stained with Ponceau S (cat#P3504; Sigma-Aldrich) to determine equivalent loading and transfer efficiency.

### Confocal imaging

*p35S:NF-YB2-YFP* and *p35S:NF-YB2^E65R^:YFP* in *nf-yb2 nf-yb3* background, and *nf-yb2 nf-yb3* seeds were cold stratified in the dark for 48-h then germinated and grown on B5 media under 24hr light. Six to seven-day-old seedlings were counterstained with propidium iodide (PI) (50μg/mL) for five minutes, washed in DI water for five minutes and whole mounted in fresh DI water on standard slides. Hypocotyls were imaged with an Olympus FluoView 500 using a 60X WLSM objective. XYZ scans were taken with line sequential scanning mode where fluorescent signals were sampled using a filter-based detection system optimized for YFP and PI with chloroplast autofluorescence also detected in the latter. YFP was excited using a 488nm Argon laser whereas PI was excited using a 543nm Helium Neon laser. Approximately 50 serial sections were imaged with a cubic voxel size of 414nm x 414nm x 414nm. Image processing took place in ImageJ (http://rsb.info.nih.gov/ij/) where average intensity projections where taken from YFP and PI channels and merged.

### Yeast two-hybrid (Y2H) and three-hybrid (Y3H) analysis

Entry clones of *NF-YA2* and *NF-YC9*, which were previously described (17, 41), were subcloned into pDEST^™^22 (Invitrogen) and pTFT1 (66) respectively to obtain an activation domain (AD) and bridge construct. The DNA binding domain (DBD) and AD constructs for *NF-YB2* and *NF-YC9* were previously described (17). The plasmids were transferred to the yeast strains MaV203 (Invitrogen) for Y2H and PJ69-4α (67) for Y3H analysis. Protein interactions were tested according to the ProQuest^™^ manual (Invitrogen). For the X-Gal assay nitrocellulose membranes were frozen in liquid nitrogen and placed on a filter paper saturated with Z-buffer containing X-Gal (5-bromo-4-chloro-3-indoxyl-beta-D-galactopyranoside, Gold Biotechnology, cat#Z4281L). For the synthetic dropout medium lacking the amino acid Histidine, 5mM 3-amino-1,2,4-triazole (3-AT) was added to eliminate nonspecific activation.

### qPCR analysis

Total RNA was collected from seven-day-old or nine-day-old seedlings according to instructions in the E.Z.N.A Plant RNA Kit (cat#R6827-01; Omega Biotek). First-strand cDNA synthesis was performed as previously described (41). For qPCR a CFX Connect^™^ Real-Time PCR Detection System (Bio-Rad) with the SYBR Green qPCR Master Mix (cat#K0222; Fermentas) was used. Results were analyzed using CFX Manager^™^ (Bio-Rad) where samples were normalized to a constitutively expressed reference gene At2G32170 (68). S1 Table lists primer sequences used for qPCR analysis.

## Author Contributions

This project was conceived by BFH, RM, CLS, NG and RWK. CLS performed all experiments except: NG performed all EMSA experiments, DSJ performed confocal imaging, ZAM performed GUS staining assays and assisted with Y2H assays. BFH, CLS, and RM wrote the manuscript and all authors read and edited the manuscript.

## Supplemental Information

**S1 Table. List of Primers**.

**S1 Fig. *NF-YA2* is expressed in *pNF-YA2:NF-YA2* plants**. Quantification of *NF-YA2* expression in *pNF-YA2:NF-YA2* plants used for qPCR analysis. Asterisks represent significant differences derived from student’s T-test (P < 0.05).

**S2 Fig. NF-YB2^E65R^ is expressed in the *nf-yb2 nf-yb3* background. (A)** Confocal images of NF-YB2 and NF-YB2^E65R^ protein localization in stable plant lines. **(B)** Protein expression in 12 individual T1 *p35S:NF-YB2^E65R^* plants compared to a stable strongly expressed *p35S:NF-YB2* in the *nf-yb2 nf-yb3* background. **(C)** Protein expression in two stable plant lines each for *p35S:NF-YB2* and *p35S:NF-YB2^E65R^* in the *nf-yb2 nf-yb3* background.

